# Caspase3-deficient cells require fibronectin for protection against autophagy-dependent death

**DOI:** 10.1101/2021.01.27.428460

**Authors:** David B. Weir, Lawrence H. Boise

## Abstract

Caspases are required for execution of apoptosis. However, in their absence, signals that typically induce apoptosis can still result in cell death. Our laboratory previously demonstrated that Casp3-deficient mouse embryonic fibroblasts (MEFs) have increased fibronectin (FN) secretion, and an adhesion-dependent survival advantage compared to wild type (WT) MEFs. Here, we show that FN is required for survival of Casp3-deficient MEFs following serum withdrawal. Furthermore, when FN is silenced, serum withdrawal-induced death is caspase-independent. However, procaspase-7 is cleaved, suggesting that MOMP is taking place. Indeed, in the absence of FN, cytochrome *c* release is increased following serum withdrawal in Casp3-deficient MEFs. Yet death does not correspond to cytochrome *c* release in Casp3-deficient MEFs. This is true both in the presence and absence of FN. Additionally, caspase-independent death is inhibited by Bcl-X_L_ overexpression. These findings suggest that Bcl-X_L_ is not inhibiting death through regulation of Bax/Bak insertion into the mitochondria, but through a different mechanism. One such possibility is autophagy and induction of autophagy is associated with caspase-independent death in Casp3-deficient cells. Importantly, when ATG5 is ablated in Casp3-deficient cells, autophagy is blocked and death is largely inhibited. Taken together, our data indicate that Casp3-deficient cells incapable of undergoing canonical serum withdrawal-induced apoptosis, are protected from autophagy-dependent death by FN-mediated adhesion.

## INTRODUCTION

Caspases are a highly conserved family of cysteine-dependent aspartate-specific proteases that are an integral part of apoptosis [1–5]. The intrinsic apoptotic pathway ultimately results in the activation of executioner caspases, at which point the enzymes complete apoptosis [6–10]. Upon a death signal, such as growth factor deprivation, Bcl-2 family proteins regulate insertion of Bax/Bak channels into the mitochondria, resulting in mitochondrial outer membrane permeabilization (MOMP) and the release of mitochondrial contents, including cytochrome *c* [11–14]. When released into the cytoplasm, cytochrome *c* triggers the formation of the apoptosome, a protein complex composed of multiple molecules each of apoptotic protease-activating factor 1 (APAF-1) and the zymogen procaspase-9. Upon recruitment to the apoptosome, the molecules of procaspase-9 become activated by proximity to one another and cleave the zymogens procaspase-3 and procaspase-7, thereby giving rise to their catalytically active forms [15–18]. Caspase-3 and caspase-7 proceed to perform internal demolition of the cell, through the selective proteolytic cleavage of hundreds of proteins [1–5, 19–21].

Our laboratory previously demonstrated that in MEFs, procaspase-3 negatively regulates fibronectin (FN) secretion, which impacts cell adhesion, migration, and survival [22]. FN is an essential component of the extracellular matrix [23–25]. Adherent cells receive prosurvival signals via integrins bound to FN and depend on integrin-fibronectin binding for survival [23, 26, 27]. Procaspase-3 regulation of FN secretion is distinct and independent from the executioner function of caspase-3 in apoptosis. Furthermore, we determined that cell survival in the absence of Casp3 occurred in an adhesion-dependent manner [22]. However, it remains unclear whether FN is necessary for this cell survival.

Here we show that FN is required for survival of Casp3-deficient cells in the absence serum. Additionally, we demonstrate that cells lacking Casp3 die following serum withdrawal through a caspase-independent mechanism when FN is absent. Furthermore, we show that this caspase-independent death involves induction of autophagy and is blocked by inhibition of autophagy. Taken together, these data indicate that when serum withdrawal-induced apoptosis is inhibited by loss of Casp3, cells become more dependent on FN-mediated adhesion that blocks an autophagy-dependent death pathway.

## RESULTS

### Fibronectin is required for survival of Casp3-deficient cells following serum withdrawal

Our previous study demonstrated that Casp3-deficient (Casp3^−/−^) mouse embryonic fibroblasts (MEFs) secrete higher levels of FN than wild type (WT) MEFs, and that their survival advantage following serum withdrawal is dependent on cell adhesion [22]. However, we never tested if this dependence is mediated by FN. Therefore, to determine the role of FN in the survival of cells lacking Casp3, we examined the effect of FN silencing on serum withdrawal-induced death in WT and Casp3^−/−^ MEFs. Specifically, we employed a doxycycline (Dox)-inducible shRNA expression system (shFN) to knockdown FN, as previously described [22] followed by serum withdrawal. A schematic of the experimental design is provided in figure 1A. Dox-induced FN knockdown was confirmed by western blot in both WT and Casp3^−/−^ (Fig. 1B). WT pLKO MEFs (control cells containing empty vector) exhibited baseline cell death of 5.31%±1.13% when seeded in medium containing serum (Fig. 1C). Death significantly increased following serum withdrawal (32.52%±2.47%) (Fig. 1C). Serum withdrawal-induced death was partially inhibited when exogenous FN was supplied (22.12%±1.87%) (Fig. 1C). Dox administration had no effect on serum withdrawal-induced death in WT pLKO MEFs in the absence or presence of exogenous FN (Fig. 1C). Importantly, serum withdrawal-induced death was nearly identical in WT shFN MEFs when Dox was not administered, consistent with a tightly regulated inducible system. However, Dox-induced FN silencing led to significantly increased death due to serum withdrawal in WT shFN MEFs (48.36%±2.22%) (Fig. 1C). This change is due to an on-target effect, as the death phenotype was significantly inhibited by exogenous FN (Fig. 1C). Casp3^−/−^ pLKO MEFs displayed baseline death of 2.22%±0.22% when cultured in medium containing serum (Fig. 1D). Consistent with our previous findings with Casp3^−/−^ MEFs [22, 28], Casp3^−/−^ pLKO MEFs are more resistant to serum withdrawal-induced death than their WT counterparts, and addition of exogenous FN resulted in a minimal, yet statistically significant decrease in death (Fig. 1D). Administration of Dox had no effect on cell death in Casp3^−/−^ pLKO MEFs, and again in the absence of Dox, shFN had no effect. However, silencing resulted in a significant increase in serum withdrawal-induced death for Casp3^−/−^ shFN MEFs, reaching 26.62%±0.97% (Fig. 1D). Thus, FN knockdown ablates the resistance of Casp3^−/−^ MEFs to serum withdrawal. Importantly, when FN was knocked down and exogenous FN was supplied, the protection from cell death seen in Casp3^−/−^ MEFs was restored, with death reduced to baseline serum withdrawal levels (Fig. 1D). Therefore, FN is required for the protection from serum withdrawal seen in Casp3-deficient MEFs.

**Fig. 1.**
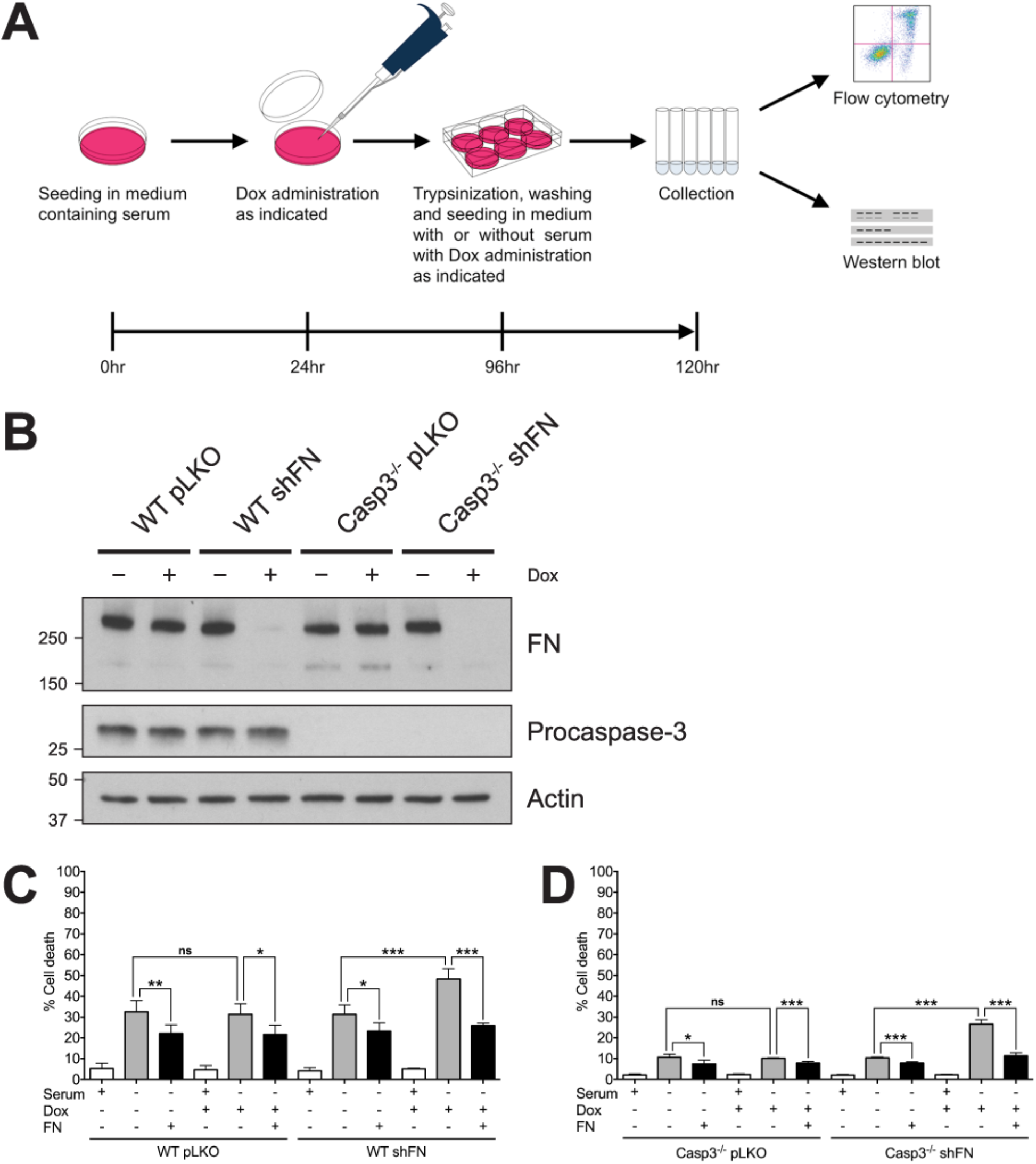
FN is required for the survival of Casp3-deficient MEFs following serum withdrawal. (A) Schematic of experimental design. (B) Cell lysates were analyzed via western blot. (C) WT pLKO and WT shFN, and (D) Casp3^−/−^ pLKO and Casp3^−/−^ shFN MEFs were seeded in medium with or without serum, in the presence or absence of Dox (5μg/ml), and incubated for 24 hr. Plates were coated with exogenous FN as indicated. Cell death was analyzed via annexin V-PI staining and flow cytometry, and was defined as annexin V+, PI+, or annexin V+/PI+. Data are represented as mean ± s.e.m. All data are from at least three independent experiments. *P < 0.05, **P = 0.01, ***P < 0.001.

### In the absence of fibronectin, serum withdrawal-induced death is caspase-independent in Casp3-deficient MEFs

Since we determined that cell survival in the absence of Casp3 is dependent upon FN, we wanted to further delineate the mechanism of cell death and whether this cell death is caspase-dependent. To this end, we included the pan-caspase inhibitor Q-VD-OPh (10μM) or a DMSO control in 24hr serum withdrawal experiments (Fig. 2). WT and Casp3^−/−^ MEFs containing the Dox-inducible shFN were seeded in media with or without serum, in the absence or presence of Dox, and in the absence or presence of Q-VD-OPh. WT shFN MEFs displayed baseline death of 5.41%±1.2% when grown in the presence of serum (Fig. 2A). Death rose to 41.20%±1.79% following serum withdrawal (Fig. 2A top panel). Death due to serum withdrawal was blocked by Q-VD-OPh, dropping to 23.93%±2.72% (Fig. 2A). In agreement with previous findings, serum withdrawal-induced cell death increased to 62.60%±4.81% when FN was knocked down. Death decreased to 29.60%±4.45% with the addition of Q-VD-OPh (Fig 2A). These data are consistent with a role for caspases in serum withdrawal-induced apoptosis. In contrast, caspase inhibition had no effect on death due to serum withdrawal in Casp3^−/−^ MEFs both with and without endogenous FN (Fig. 2B). Casp3^−/−^ MEFs grown in medium containing serum exhibited baseline death of 4.89%±1.33% (Fig. 2B). Death increased to 16.97%±01.02% following serum withdrawal, and was unaffected by addition of Q-VD-OPh (Fig. 2B). In the absence of FN, cell death was increased following serum withdrawal. However, in contrast to WT cells this was unaffected by Q-VD-OPh, suggesting that death is caspase-independent. To assure caspase activity was inhibited, we performed western blot analysis of caspase-3 and caspase-7 in the absence and presence of Q-VD-OPh (Fig. 2C). As expected, in WT shFN MEFs that were not administered Dox, cleavage of both procaspase-3 and -7 was evident under serum withdrawal (Fig. 2C left panel). Cleavage of both procaspase-3 and -7 was significantly inhibited when Q-VD-OPh was added under these conditions (Fig. 2C left panel). The amount of procaspase-3 and -7 cleavage increased when FN was knocked down, as compared to control (Fig. 2C left panel). This cleavage was also significantly inhibited in the presence Q-VD-OPh (Fig. 2C left panel). Casp3^−/−^ shFN MEFs displayed procaspase-7 cleavage when serum starved, and cleavage was reduced by Q-VD-OPh administration (Fig. 2C right panel). In contrast to WT shFN MEFs, Casp3^−/−^ shFN MEFs exhibited the same amount of procaspase-7 cleavage when FN was knocked down as when it was at baseline. Cleavage was inhibited by Q-VD-OPh when FN was knocked down as it was when FN was at baseline (Fig. 2C right panel). These data indicate that Casp3^−/−^ MEFs subjected to FN knockdown and serum withdrawal die in a caspase-independent manner. They are also consistent with our previous findings that caspase-7 is dispensable for serum withdrawal-induced death [22, 28].

**Fig. 2.**
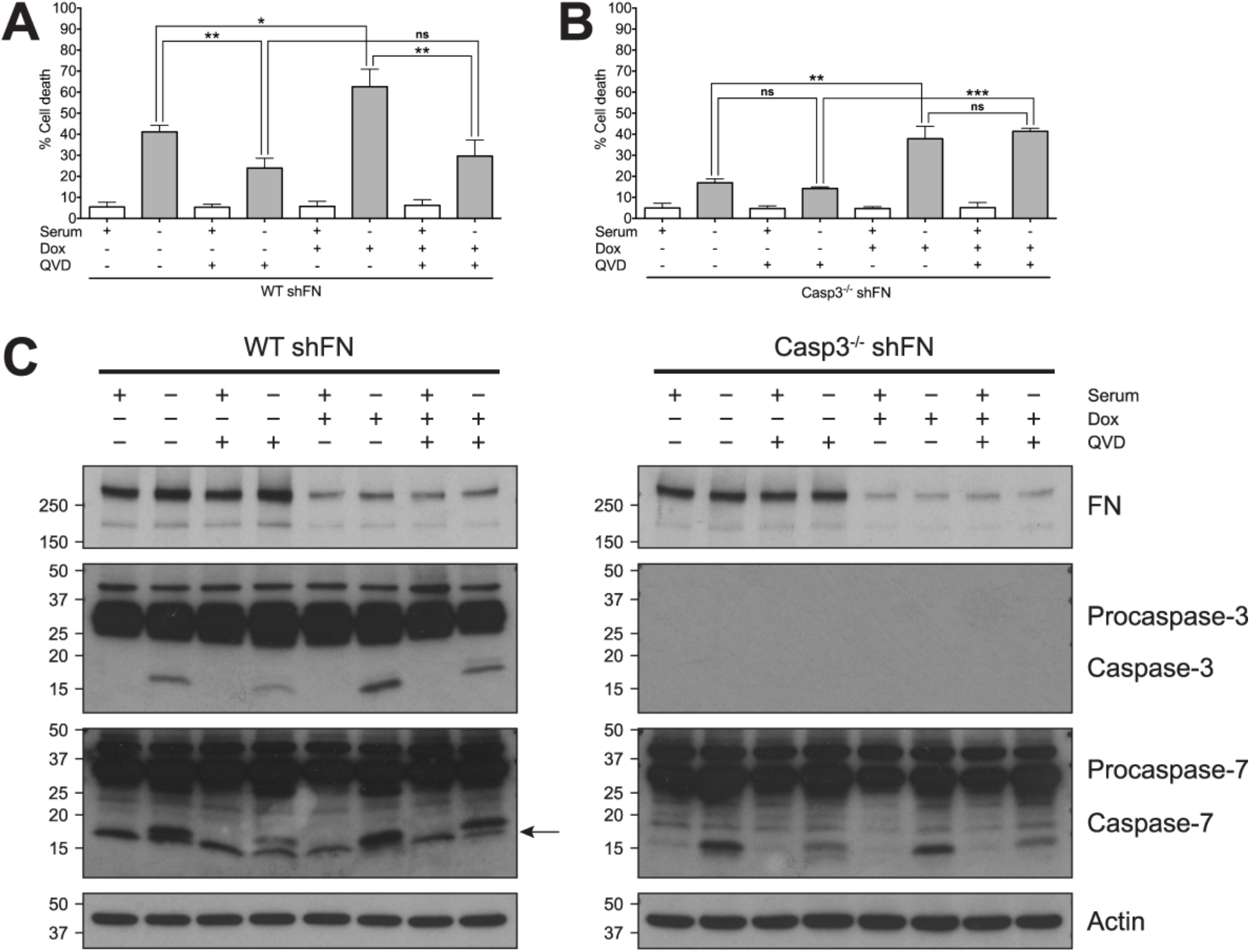
In the absence of FN, serum withdrawal results in caspase-independent cell death in Casp3-deficient MEFs. (A) WT shFN and (B) Casp3^−/−^ shFN MEFs were seeded in medium with or without serum, in the absence or presence of Dox (5μg/ml), in the absence or presence of Q-VD-OPh (QVD, 10μM), and incubated for 24hr. Cell death was analyzed via annexin V-PI staining and flow cytometry, and was defined as in Figure 1. (C) Cell lysates were analyzed via western blot. Arrow denotes a non-specific band. Data are represented as mean ± s.e.m. All data are from at least three independent experiments. *P = 0.014, **P < 0.01, ***P < 0.0001.

### Caspase-independent death following serum and fibronectin withdrawal results in the induction of MOMP and is inhibited by Bcl-X_L_

Cleavage of procaspase-7 in Casp3-deficient MEFs subjected to serum withdrawal when FN is silenced implies that mitochondrial outer membrane permeabilization (MOMP) is occurring. To explore this possibility, we examined how MOMP inhibition affects caspase-independent cell death by overexpressing Bcl-X_L_, a prosurvival member of the Bcl-2 protein family, in WT and Casp3-deficient MEFs. Cell lines containing a Bcl-X_L_ expression vector or empty vector were generated (Fig. 3A) and seeded into media with or without serum, in the absence or presence of Dox (Fig. 3B,C,D). To measure the degree of MOMP in each set of conditions, we employed a flow cytometric cytochrome *c* release assay [29]. Representative flow cytometric plots are included in figure 3B. Briefly, following permeabilization, staining, and fixing, intact cells were sorted by light scatter characteristics, and changes in cytochrome *c* staining intensity were detected. Both serum withdrawal-induced cell death and cytochrome *c* release were measured in vector control (pBABE) cells (Fig. 3C) and Bcl-X_L_ overexpressing cells (Fig. 3D). The antibiotic alamethicin was used as a positive control for cytochrome *c* release. Under all conditions, WT shFN pBABE cells displayed corresponding death and cytochrome *c* release. These data are consistent with activation of the intrinsic apoptotic pathway following serum withdrawal with inhibition of FN secretion resulting in increased intrinsic apoptotic signaling. In contrast, in Casp3^−/−^ cells, death did not mirror cytochrome *c* release. In fact, there was no difference between cytochrome *c* release in WT and Casp3^−/−^ cells, while significant differences were observed in cell death. This suggests that all of the differences between the WT and Casp3^−/−^ cells occurs downstream of MOMP, a finding consistent with loss of an effector caspase. However, when Bcl-X_L_ was introduced into the cells, it was able to block both cytochrome *c* release and cell death regardless of caspase-3 expression (Fig. 3D). Together, these findings suggest that in the absence of caspase-3, the caspase-independent death that is observed is inhibitable by Bcl-X_L_. One possibility is that the death is due to MOMP itself. However, when FN is present, MOMP levels were high yet death was low (Fig. 3C). Therefore, we investigated the role of an alternate Bcl-X_L_-inhibitable pathway, autophagy.

**Fig. 3.**
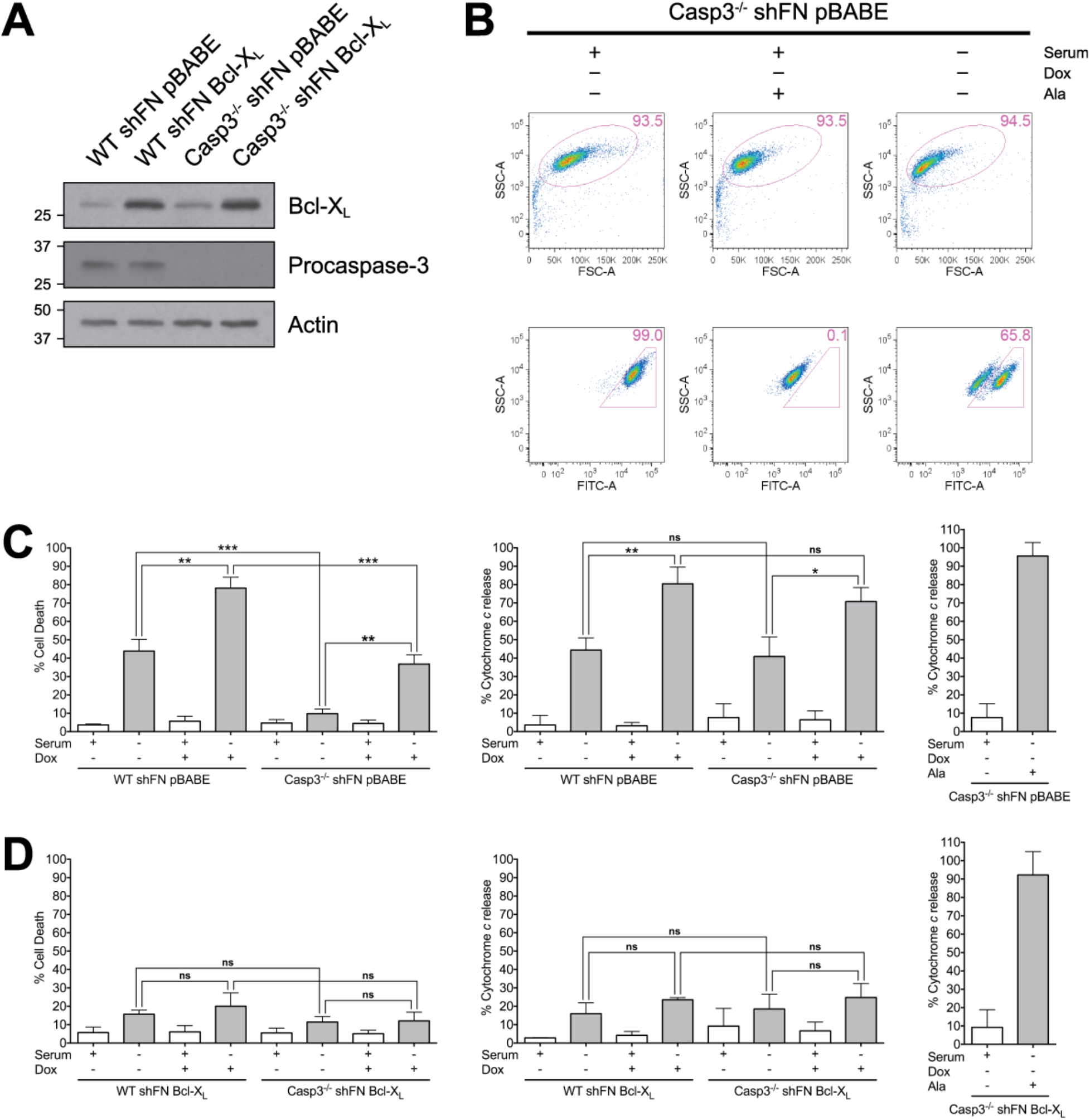
Cytochrome *c* release is independent of serum withdrawal-induced death in Casp3-deficient MEFs. (A) WT shFN and Casp3^−/−^ shFN MEF cell lines with empty expression vector pBABE-hygro (pBABE) or vector containing Bcl-X_L_ were generated (WT shFN pBABE, WT shFN Bcl-X_L_, C3^−/−^ shFN pBABE, and C3^−/−^ shFN Bcl-X_L_). Cell lysates were analyzed via western blot. (B) Representative flow cytometry plots from cytochrome *c* release assay. (C, left panel) WT shFN pBABE and C3^−/−^ shFN pBABE, and (D, left panel) WT shFN Bcl-X_L_ and C3^−/−^ shFN Bcl-X_L_ MEFs were seeded in medium with or without serum, in the absence or presence of Dox (5μg/ml), and incubated for 24hr. Cell death was analyzed via annexin V-PI staining and flow cytometry, and was defined as in Figure 1. (C center,right panels) WT shFN pBABE and C3^−/−^ shFN pBABE, and (D center,right panels) WT shFN Bcl-X_L_ and C3^−/−^ shFN Bcl-X_L_ MEFs were permeabilized in the absence or presence of alamethicin, stained with a cytochrome *c* antibody conjugated to Alexa Fluor 488, and analyzed via flow cytometry. Data are represented as mean ± s.e.m. All data are from at least three independent experiments. *P = 0.0168, **P < 0.01, ***P ≤ 0.001.

### Induction of autophagy is associated with death in Casp3-deficient cells

To determine whether autophagy is occurring in Casp-3-deficient MEFs subjected to FN knockdown and serum withdrawal, we measured lipidated-LC3B (LC3B-II) via western blot (Fig. 4). Serum withdrawal resulted in an increase in LC3B-II in both WT (Fig. 4A,C) and Casp3^−/−^ (Fig. 4B,D) cells and this was enhanced by FN silencing. Additionally, LC3B-II accumulation was blocked by Bcl-X_L_ overexpression. To assure that LC3B-II induction was due to an increase in autophagy and not a decrease in autophagic flux, we repeated the experiment in Casp3-deficient cells (Casp3^−/−^ shFN pBABE LCv2 MEFs) and added bafilomycin A1 (Baf) (100nM) or a DMSO control for the final 4 hours to block autophagosomal protein degradation. Baf addition resulted in further increase in LC3B-II (Fig. S1A,B), indicating that the serum withdrawal-induced changes were due to an increase in autophagy. LC3B-II was not observed in cells lacking ATG5 (Casp3^−/−^ shFN ATG5^−/−^ 1 MEFs) under any conditions tested (Fig. S1A). Baf administration had no effect on cell death in serum withdrawal experiments (Fig. S1C). We next quantified the change in LC3B-II as per recommended methods (see Materials and Methods) across several experiments. Interestingly, the increase in LC3B-II, and by inference the induction of autophagy, was greater in the Casp3-deficient cells following serum withdrawal both in the presence and absence of endogenous FN (Fig. 4). This is consistent with the concept that autophagy is limited by the induction of apoptosis and suggests that autophagy is unchecked in the Casp3-deficient MEFs, resulting in a 40-fold increase when FN was silenced and serum was removed. This raised the possibility that the caspase-independent death that occurs under these conditions is due to excess autophagy.

**Fig. 4.**
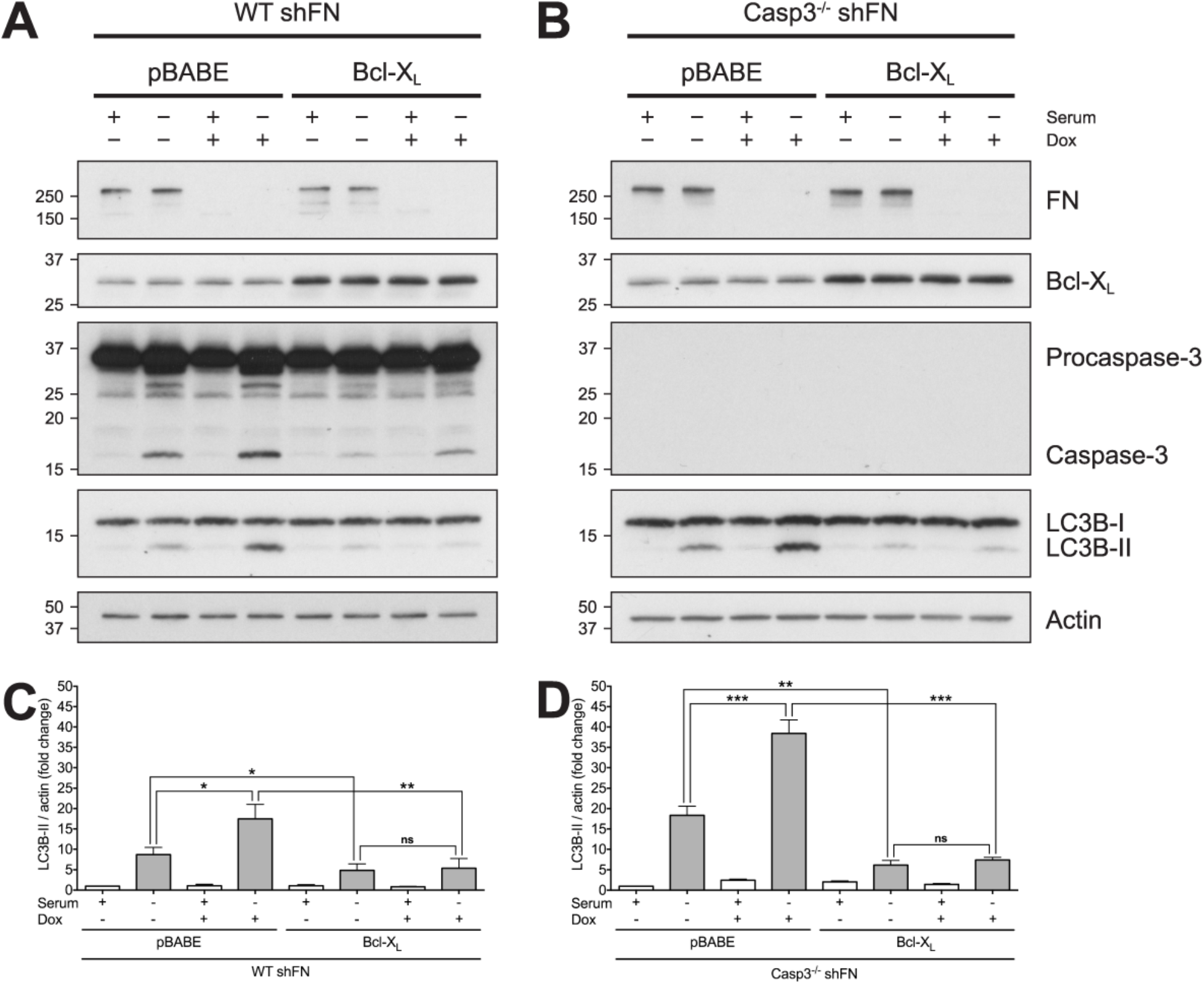
Autophagy is increased by serum withdrawal in Casp3-deficient MEFs when FN is silenced. Cell lysates from subsets of (A) WT shFN pBABE and WT shFN Bcl-X_L_, and (B) Casp3^−/−^ shFN pBABE MEFs and C3^−/−^ shFN Bcl-X_L_ MEFs in Figure 3 were analyzed via western blot. (C,D) Quantification of LC3B-II in panels A and B, respectively. Data are represented as mean ± s.e.m. All data are from at least three independent experiments. *P < 0.05, **P < 0.01, ***P ≤ 0.001.

### Autophagy inhibition blocks death in Casp3-deficient cells

Since we had demonstrated that both cell death and the amount of autophagy induced by serum withdrawal in Casp-3-deficient MEFs increase significantly when FN is knocked down, we investigated whether autophagy is required for cell death under conditions of FN knockdown and serum withdrawal. In order to test this, we used CRISPR-Cas9 to knock out ATG5. A single cell clone containing the empty vector (Casp3^−/−^ shFN pBABE LCv2 MEFs) was used as a control. Additionally, a single cell clone that received the vector containing a single guide RNA (sgRNA) but retained ATG5 expression (Casp3^−/−^ shFN pBABE ATG5^+/+^) was used as an additional control. Two single cell clones that received the same sgRNA as the Casp3^−/−^ shFN pBABE ATG5^+/+^ MEFs but lost ATG5 (Casp3^−/−^ shFN pBABE ATG5^−/−^ 1 and Casp3^−/−^ shFN pBABE ATG5^−/−^ 2 MEFs) were tested to determine the effect of ATG5 knockout on serum withdrawal with and without endogenous FN. Consistent with Figure 4, following serum withdrawal, Casp3-deficient MEFs that retained ATG5 showed patterns of LC3B-II consistent with induction of autophagy (Fig. 5B,C). In contrast, in both ATG5-deficient clones, serum withdrawal-induced autophagy was completely blocked as evidenced by a complete block in LC3B lipidation (Fig. 5B). Importantly, ATG5 loss and autophagy induction resulted in a complete block of cell death in FN-silenced cells following serum withdrawal (Fig. 5D). To ensure that loss of the autophagic function, and not an alternate function, of ATG5 is responsible for blockade of cell death, we separately used CRISPR-Cas9 to knock out a second gene required by the autophagy pathway, Autophagy Related Gene 7 (ATG7). A population of cells that grew out following infection and selection as a bulk culture containing the empty vector (Casp3^−/−^ shFN pBABE LCv2 Bulk MEFs) was used as a control, and a similarly selected population of ATG7 knockout cells (Casp3^−/−^ shFN pBABE ATG7^−/−^) were tested. Following serum withdrawal, Casp3-deficient MEFs expressing ATG7 displayed LC3B-II levels corresponding to autophagy induction (Fig. S2A,B). Conversely, in ATG7 knockout cells, serum withdrawal-induced autophagy was almost completely blocked, as indicated by a nearly total loss of LC3B lipidation (Fig. S2A,B). In agreement with results from ATG5 knockout experiments, death in FN-silenced cells following serum withdrawal was inhibited by ATG7 knockout (Fig. S2C). Taken together, these data indicate that death due to serum withdrawal in Casp3-deficient MEFs with diminished FN secretion requires autophagy.

**Fig. 5.**
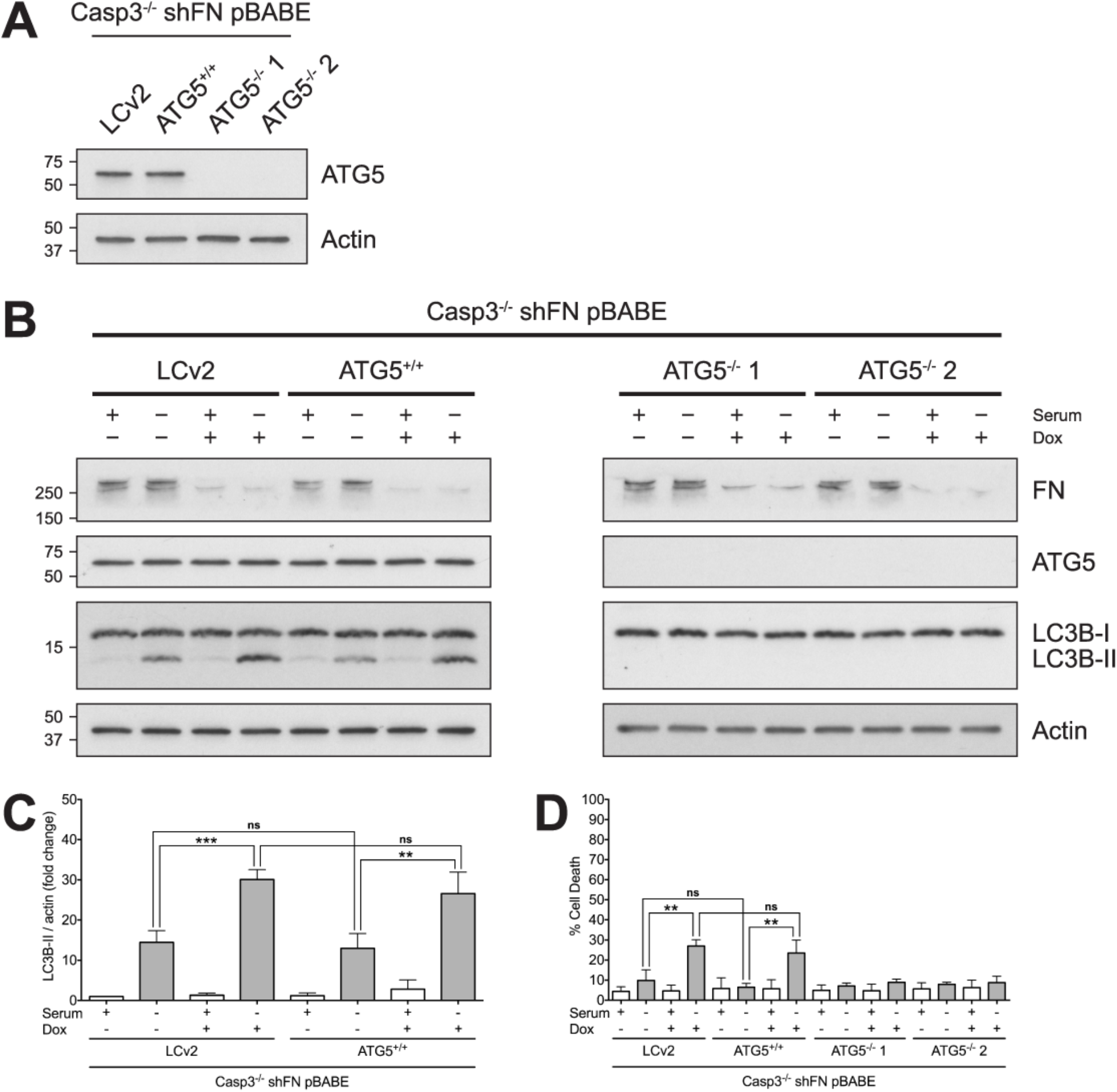
Ablation of ATG5 expression blocks serum withdrawal-induced death in Casp3-deficient MEFs. (A) Single cell clones with functional ATG5 (Casp3^−/−^ shFN pBABE LCv2 and Casp3^−/−^ shFN pBABE ATG5^+/+^) and single cell clones with loss of ATG5 expression (Casp3^−/−^ shFN pBABE ATG5^−/−^ 1 and Casp3^−/−^ shFN pBABE ATG5^−/−^ 2) were generated. Cell lysates were analyzed via western blot. (B) Casp3^−/−^ shFN pBABE LCv2, C3^−/−^ shFN pBABE ATG5^+/+^, Casp3^−/−^ shFN pBABE ATG5^−/−^ 1, and Casp3^−/−^ shFN pBABE ATG5^−/−^ 2 MEFs were seeded in medium with or without serum, in the absence or presence of Dox (5μg/ml), and incubated for 24hr. Cell lysates were analyzed via western blot. (C) Quantification of LC3B-II in Casp3^−/−^ shFN pBABE LCv2 and C3^−/−^ shFN pBABE ATG5^+/+^ MEFs in panel B. (D) Cell death was analyzed via annexin V-PI staining and flow cytometry, and was defined as in Figure 1. Data are represented as mean ± s.e.m. All data are from at least three independent experiments. **P < 0.01, ***P = 0.0002.

## DISCUSSION

Apoptosis and autophagy have a complex connection [30–33]. The two pathways share molecular components, can be triggered by the same stimuli, and are capable of interacting in several distinct ways [30–34]. They can work in collaboration [35–38], as well as in opposition to one another [39–44]. Additionally, although autophagy most often serves as a cytoprotective process [45–47], it can also function as a killing mechanism under certain conditions, including when a cell is incapable of performing apoptosis [48–57]. Bax/Bak-deficient MEFs were found to undergo a non-apoptotic cell death program that is dependent upon autophagic proteins when treated with etoposide [48]. Additionally, bone marrow-derived IL-3-dependent Bax/Bak-deficient mouse cells activate autophagy and ultimately succumb to cell death, when withdrawn from IL-3 [49]. Furthermore, zVAD was demonstrated to induce autophagic cell death in L929 cells when caspase-8 expression was decreased [53]. In this study, we demonstrate that cells lacking Casp3 become dependent on FN for survival following serum withdrawal, and cells undergo non-apoptotic, autophagy-dependent death.

Our laboratory previously showed that procaspase-3 is a negative regulator of FN secretion and impacts adhesion, migration and survival independent of catalytic function [22]. Introduction of WT or a catalytically inactive mutant caspase-3 reversed the adhesion and migration phenotypes seen in Casp3 deficient MEFs [22]. Additionally, we showed that casp3-deficient MEFs have an adhesion-dependent survival advantage over WT MEFs following serum withdrawal [22]. However, introduction of a catalytically inactive mutant caspase-3, had no effect on survival of Casp3 deficient MEFs following serum withdrawal [58]. These data indicate that loss of caspase-3, and not the effects on increased FN secretion were primarily responsible for increased cell survival. The change in FN secretion due to procaspase-3 loss was only a two-fold change [22]. In the current study we investigated the effect of loss of FN on the survival of Casp3-deficient MEFs. Here, we show that in Casp3-deficient MEFs, FN is required for protection from caspase-independent death following serum withdrawal. These findings agree with previous data showing that caspase-7 is dispensable for this death mechanism [28]. Following serum withdrawal, autophagy is induced in Casp3-deficient cells, and cells survive. Interestingly, when FN is silenced, autophagy increases significantly, and death occurs. This raises the possibility that death is contingent upon autophagy. Importantly, blocking autophagy inhibits serum withdrawal-induced death in Casp3-deficient cells, indicating that death is indeed autophagy-dependent.

In WT MEFs, serum withdrawal triggers MOMP, as well as autophagy signaling. MOMP leads to activation of caspase-3, which in turn both inhibits autophagy and completes apoptosis. Inhibition of autophagy by caspase-3 is well documented [39, 40, 59]. Therefore, it is reasonable to postulate that loss of caspase-3 not only blocks apoptosis, but releases constraints on autophagy. Our model (Fig. 6) posits that serum withdrawal of Casp3-deficient MEFs stimulates MOMP and induces an increased amount of autophagy. However, these cells survive serum withdrawal, primarily because they cannot complete canonical apoptosis due to their lack of caspase-3. Importantly, in the absence of FN, serum withdrawal induces still more autophagy, and under these conditions cell death occurs. Thus, in Casp3-deficient cells, FN blocks death that results if autophagy proceeds unhampered.

**Fig. 6.**
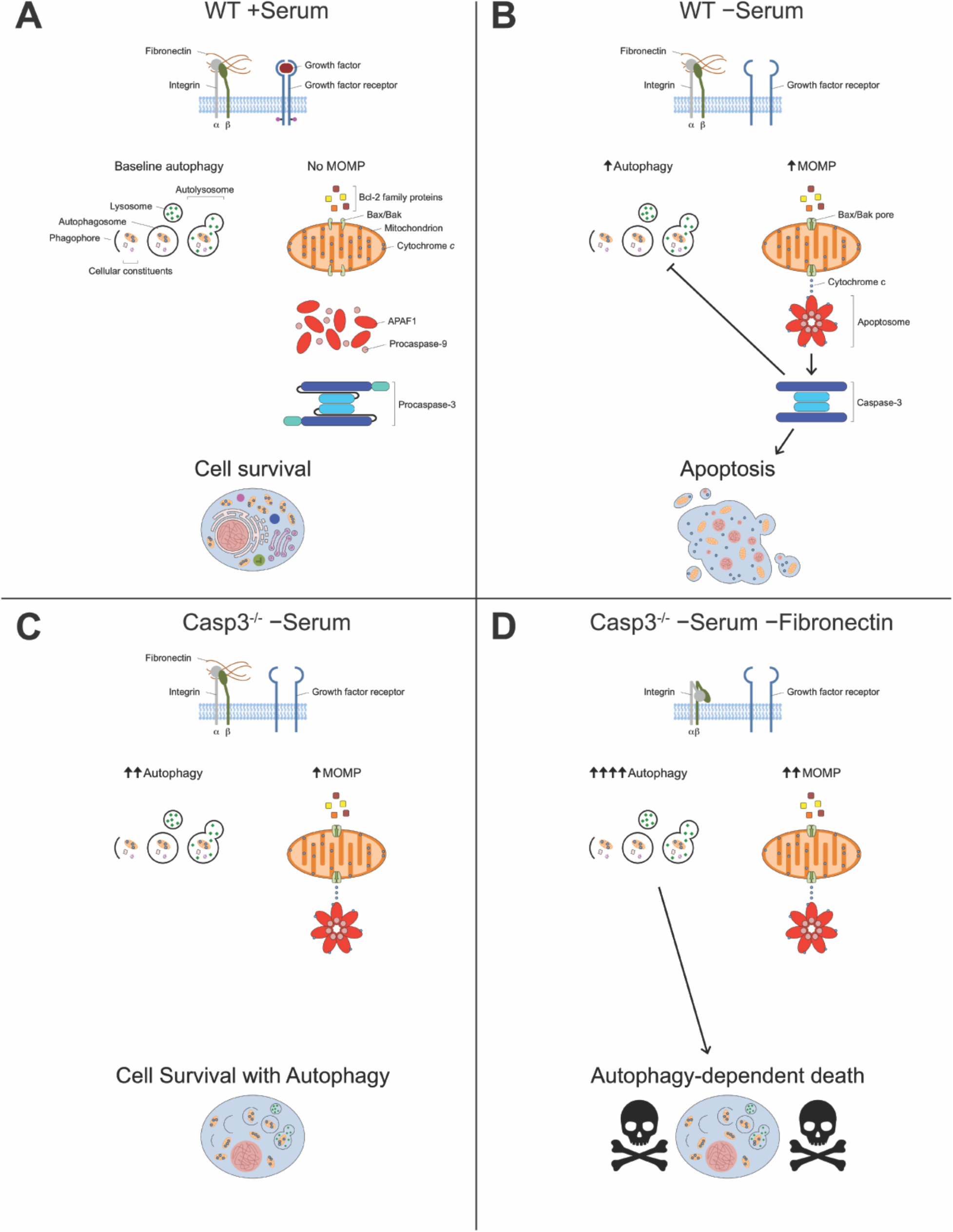
Casp3-deficient MEFs require FN for protection against autophagy-dependent death following serum withdrawal. (A) WT MEFs grown in medium containing serum have both FN and growth factors for integrins and growth factor receptors to bind to, respectively. Under these conditions, cells have baseline autophagy and no MOMP takes place. (B) Serum withdrawal induces both autophagy and MOMP in WT MEFs. MOMP allows release of cytochrome *c* from mitochondria, which leads to activation of caspase-3. Once activated, caspase-3 both inhibits autophagy and completes apoptosis. (C) In Casp3-deficient MEFs, serum withdrawal induces increased autophagy compared to WT MEFs, but the same amount of MOMP. Caspase-3 is not present to inhibit autophagy or complete apoptosis, and cells survive. (D) In the absence of FN, serum withdrawal induces still more autophagy, as well as increased MOMP in Casp3-deficient MEFs. Cells die in an autophagy-dependent, non-apoptotic fashion.

These findings open up several important questions. For example, how exactly are Casp3-deficient cells in the absence of FN dying following serum withdrawal? Cells may be undergoing autophagy-dependent cell death, but not death directly caused by autophagy itself. Autophagy may be activating another death pathway that ultimately kills the cell. Evidently, cells are not dying via necroptosis, as death was not blocked by RIPK3 inhibition (data not shown). Furthermore, a lack of membrane disruption is observed under these experimental conditions (Fig. S3). Although we have not ruled out ferroptosis or pyroptosis, based on this finding, it is unlikely that either is involved.

Additionally, how does FN protect Casp3-deficient cells from death? Perhaps Casp3-deficient cells are resistant to loss of one prosurvival signal, i.e., serum or FN signaling, but are not capable of withstanding the loss of both signals. Excessive autophagy may result when both survival signals are lost. Alternatively, FN signaling via integrins may modulate the cellular response to serum withdrawal. Crosstalk between integrins and growth factor receptors are important signaling mechanisms [60–62]. Coupling integrin signaling and growth factor receptor signaling is crucial for cultivating the appropriate cellular response for a given context [60–62]. The autophagy signal received by cells due to the absence of growth factors may be dampened by FN signaling. Possible signaling pathways involved include Ras-Raf-MEK-ERK and PI3K/Akt. In contrast, the level of autophagy induced by serum withdrawal in the absence of FN may be the same as in its presence, but the signal is never turned off, resulting in excessive autophagy. Furthermore, although the prosurvival signal arising from FN is almost definitely mediated by integrins, it is not clear which integrins are involved. At least a dozen integrin dimers bind FN [23]. We have previously shown that Casp3-deficient MEFs have cell surface expression of integrins β_1_ and β_3_ [22]. Whether an individual integrin dimer or a combination of integrin dimers binds FN under experimental conditions is yet to be determined.

Taken together, our data demonstrate that loss of Casp3 results in cellular dependence on FN for protection from autophagy-dependent death. This may have implications in disease settings, such as cancer, where adhesion, autophagy and cell death all play crucial roles. It would be expected that cancer cells would lose caspase-3, as a means of evading apoptosis, yet this is rarely the case [63]. In fact, caspase-3 expression has been found to be elevated in a number of cancer types [64–66]. Perhaps losing caspase-3 would leave cells vulnerable to excessive autophagy. Autophagy has complicated and sometimes diametrically opposed roles in cancer [41, 42, 67–69]. A number of studies suggest autophagy serves as a barrier against oncogenesis [70–74]. Conversely, compelling evidence supports a role for autophagy in survival of established cancer cells [42, 67, 68, 75–77]. Genes encoding central components of the autophagy pathway are generally not lost or mutated in cancer [78–80]. However, these types of mutations do occur in specific cancer types [80–82]. Heterozygous deletion of beclin 1 is found in 40-75% of sporadic breast, ovarian and prostate cancer [81, 82]. Three core autophagy genes, RB1CC1/FIP200, ULK4, and WDR45/WIPI4, were found to be under positive selection for somatic mutations in endometrial carcinoma [80]. This is also true of ATG7 in clear cell renal carcinoma [80]. Frameshift mutations in ATG2B, ATG5, ATG9B, and ATG12 were found in more than a quarter of gastric and colorectal cancers with microsatellite instability [83]. The significance of these recurrent mutations in certain cancer types is yet to be determined. Perhaps restricting autophagy in tumor cells where autophagy is already very high provides protection from death. It has been suggested that autophagy might cause cancer cell death [84]. Our data suggests loss of caspase-3 might increase autophagy in cancer cells even further, potentially leaving them susceptible to loss of additional survival signaling, such as cell adhesion. If such signaling were to decrease, cancer cells could undergo autophagy-dependent cell death. Interestingly, this may also point to an additional reason that gains in anti-apoptotic Bcl-2 proteins are observed in cancer. In contrast to loss of caspase-3, gain of Bcl-2 blocks both apoptosis and autophagy, and can overcome this internal checkpoint for induction of cell death by multiple mechanisms. A number of cancer therapies induce autophagy [85–90]. When administered low doses of estrogen agonists, MCF-7 breast cancer cells, one of the few cancer cells lacking caspase-3, were shown to have extensive autophagosome formation and die by a nonapoptotic mechanism [90]. Promoting autophagy has been suggested as a therapeutic approach to cancer [69, 91]. Indeed, numerous clinical trials have investigated cancer therapies that induce autophagy [92–95]. However, most of these trials do not involve intentional enhancement of autophagy and do not attempt to determine if autophagy is increased by treatment [96]. The vast majority of previous investigation focused on autophagy-based therapies has involved inhibiting it [69, 91]. Our findings provide evidence to suggest that research focused on autophagy-based cancer therapies that enhance autophagy may be further warranted.

## MATERIALS AND METHODS

### Cell culture

MEFs, ΦNX-ecotropic, and HEK293T (American Type Culture Collection) cells were cultured and split as previously described [22].

### Retroviral transduction and generation of stable cell lines

ΦNX-ecotropic packaging cells were transfected with pBABE-hygro (Addgene plasmid # 1765; empty or Bcl-X_L_) using Lipofectamine 2000 (Invitrogen). Target MEFs were retrovirally transduced as previously described [22]. Once cells recovered from infection, they were selected through hygromycin B (Calbiochem) selection.

### Lentiviral transduction and generation of stable cell lines

HEK293T packaging cells were co-transfected with lentiCRISPR v2-Blast (Addgene plasmid # 83480; empty, ATG5-specific sgRNA [AAGATGTGCTTCGAGATGTG; designed using crispr.mit.edu], or ATG7-specific sgRNA [GAACGAGTACCGCCTGGACG; designed using crispor.tefor.net]), psPAX2 (Addgene plasmid # 12260), and pMD2.G (Addgene plasmid # 12259) using jetPRIME transfection reagent (Polyplus) according to the manufacturer’s instructions. Fresh medium was added, and cells were incubated for 48hr at 37°C in a humid incubator under 5% CO_2_. Target MEFs were seeded on six-well plates and allowed to grow for 24 hours. Viral supernatents were collected and filtered through 0.45μm syringe filters (Pall), at 48 hours. Viral supernatents were applied directly to target MEFs for infection using Polybrene Infection/Transfection Reagent (Millipore). After 24 hours, viral supernatents were removed and replaced with fresh medium. Once cells had recovered from infection, they were selected with blasticidin (Thermo Fisher).

### Western blot analysis

SDS/PAGE and western blot analysis were performed as previously described [97]. Primary antibodies against Caspase-3 (Cell Signaling), Caspase-7 (Cell Signaling), Fibronectin (Abcam), Bcl-X_L_ (Cell Signaling), LC3B (Sigma), ATG5 (Cell Signaling) and actin (Sigma) were used.

### FN knockdown and coating

Knockdown of FN was performed as previously described [22]. FN coating was accomplished by adding phosphate buffered saline (PBS, Cellgro) and human plasma fibronectin (Millipore) to six-well plates (Corning). Plates were incubated for 4 hours at 37°C in a humid incubator under 5% CO_2_. PBS was removed immediately prior to experiments.

### Cell death assays

MEFs were grown for 24 hours and subsequently administered doxycycline (Sigma) for 72 hours, as indicated. Cells were then trypsinized for 5 minutes. Trypsin was inactivated using medium containing serum. Cells were then washed three times with serum-free medium, seeded into six-well plates and treated as indicated. After 24 hours, the medium and adherent cells were harvested, washed with PBS, and stained with annexin V-FITC (BioVision) and propidium iodide (PI, Sigma), as described previously [98]. Cell death was determined by flow cytometry on a BD FACSCanto II system with FACSDiva software, and data was analyzed with FlowJo.

### Caspase inhibition assays

Inhibition of caspases was accomplished through administration of the pan-caspase inhibitor Q-VD-OPh (BioVision).

### Cytochrome *c* release assay

After serum withdrawal, cells were collected, subjected to digitonin treatment, fixed, stained with an Alexa Fluor 488 anti-cytochrome *c* antibody (BD), and analyzed via flow cytometry. Where indicated, cells received alamethicin treatment concurrent with digitonin treatment, and served as positive controls. Cytochrome *c* release was determined by flow cytometry on a BD FACSCanto II system with FACSDiva software, and data was analyzed with FlowJo.

### Densitometry and LC3B-II/actin quantification

LC3B-II/actin quantification and comparison were accomplished via methods widely used to monitor autophagy [99]. Briefly, Fiji software was used to calculate OD values for LC3B-II and actin for each condition of three different western blots from independent sets of lysates. The ratio of LC3B-II to actin (LC3B-II/actin) was expressed as a fold change normalized to negative control cells.

### Lysosomal acidification inhibition assay

Acidification of the lysosome was inhibited through administration of the V-ATPase inhibitor Bafilomycin A1 (Cell signaling).

### Necroptosis assay

MEFs were grown for 24 hours and subsequently administered doxycycline (Sigma) for 72 hours, as indicated. Cells were then trypsinized for 5 minutes. Trypsin was inactivated using medium containing serum. Cells were then washed three times with serum-free medium, seeded into six-well plates, and treated as indicated, including treatment with a necroptosis-inducing cocktail consisting of the SMAC mimetic BV6 (Calbiochem), TNFα (Peprotech) and zVAD-fmk (Enzo). SYTOX Green (Invitrogen) was included in all conditions as an indicator of necroptosis. Necroptosis was determined by live-cell imaging on an IncuCyte Zoom system (Essen), data was analyzed with IncuCyte Zoom software (Essen).

### Statistics

Student’s unpaired *t*-test was used to analyze data from at least three independent experiments. Data are represented as mean ± standard error of the mean.

## Acknowledgements

This work was supported by R01 GM106565 (LHB). The authors thank Edward S. Mocarski for reagents and use of the Incucyte instrument to measure necroptosis. MEFs were a gift from Richard Flavell. ΦNX-ecotropic cells were a gift from Garry Nolan. pBABE-hygro was a gift from Hartmut Land, Jay Morgenstern and Robert Weinberg. lentiCRISPR v2-Blast was a gift from Mohan Babu. psPAX2 and pMD2.G were gifts from Didier Trono. We also thank Adam Marcus and Shannon Matulis for critical review of the manuscript.

